# Inhibition of IL-22-producing cells by *Sutterella* sp. bacteria

**DOI:** 10.1101/2025.03.20.644103

**Authors:** Louise Dupraz, Giovanna Orianne, Grégory Da Costa, Romain Gauthier, Amy Blondeau, Marie Boinet, Laura Creusot, Aurélie Magniez, Loïc Chollet, Nathalie Rolhion, Camille Danne, Zdenek Dvorák, Philippe Seksik, Harry Sokol, Marie-Laure Michel

## Abstract

The gut microbiota constitutes a complex ecosystem essential for host defense against infection and maturation of the immune system. Inflammatory bowel diseases (IBD), such as ulcerative colitis and Crohn’s disease, are characterized by a severe inflammation of the intestine, arising from dysregulated control of host-microbiota crosstalk. However, neither the genetic bases of IBD nor the immune responses involved are fully understood. The pathobionts are currently under investigation for their active role in the development and the severity of IBD. These bacteria are present in the microbiota of healthy individuals without causing disease but have pathogenic potential when the intestinal environment is disturbed. Here, we highlighted *Sutterella* sp. as a new commensal pathobiont for its capacity to modulate the host immune functions. This anaerobic Gram-negative bacterium inhibits IL-22 production and AhR activity in mice and humans. The effectors responsible for the biological activity are large (>30 kDa) protein-based compounds secreted by *Sutterella* sp. These data enhance our understanding of the mechanisms that regulate host immune functions and pave the way for new therapeutic strategies to control gut inflammation.

## INTRODUCTION

The gut microbiota forms a complex ecosystem that plays a crucial role in host defense against infections and the maturation of the immune system^1^. Inflammatory bowel diseases (IBD), such as ulcerative colitis (UC) and Crohn’s disease (CD), are characterized by severe intestinal inflammation resulting from dysregulated interactions between the host and its microbiota^1^. These pathologies require long-term treatments based on anti-inflammatory and immunosuppressive therapies, which are often associated with drug-related side effects. Moreover, some patients exhibit poor responses to existing therapies, highlighting the need for novel treatment approaches. Modulation of the gut microbiota is an actively studied strategy, notably through the use of probiotics, prebiotics, or fecal transplantation to enhance beneficial microbial species (such as *Faecalibacterium prausnitzii*) or microbiota-derived metabolites (such as short-chain fatty acids and aryl hydrocarbon receptor (AhR) ligands) that support gut health^2^.

The pathobionts are also currently under investigation for their active role in the development and severity of IBD^3^. These are bacteria present in the microbiota of healthy individuals without causing disease but they can become pathogenic when the intestinal environment is disturbed. Research has primarily focused on adherent invasive *Escherichia coli* (AIEC), *Helicobacter hepaticus* or *Ruminococcus gnavus*^*3*^. Identifying additional pathobionts and deciphering their mechanisms of action could lead to a better understanding and management of IBD pathophysiology and pave the way for a new generation of treatment.

Interleukin-22 (IL-22) is a member of IL-10 family, crucial to maintain epithelial barrier function, promote tissue repair and protect against pathogens and pathobionts^4, 5^. This cytokine is produced by αβ CD4^+^ T helper 17 (Th17) cells, T helper 22 (Th22) cells, γδ T cells and ROR-γt^+^ innate lymphoid cells (ILC3 CD4^+^ and CD4^neg^), in response to IL-23 and IL-1β produced by dendritic cells and macrophages. IL-22 targets mainly tissue epithelial cells that express IL-22R1, stimulates the production of protective mucus from goblet cells, and induces the production of antimicrobial peptides (RegIIIβ, RegIIIγ and β-defensins)^6^. IL-22 protects the intestine against bacterial pathogens such as *Citrobacter rodentium*^*7*^ *and Clostridium difficile*^*8*^, but favors *Salmonella typhimurium* colonization^9^ and *Toxoplasma gondii*-induced immunopathology^10^. During intestinal inflammation, IL-22 is present in high quantities in the blood of patients with Crohn’s disease (CD). It protects intestinal mucosa from inflammation *via* the maintenance of epithelial barrier integrity in CD^11^. It also has a protective effect in DSS (dextran sulfate sodium)-induced colitis mouse model^12^, since genetic or pharmacological ablation of IL-22 induces an exacerbated epithelial destruction and colonic inflammation. However, IL-22 also worsens inflammation in a chronic colitis model that is induced by an adoptive transfer of memory CD4^+^ T cells^13^. IL-22 was thus shown to play beneficial or deleterious roles in intestinal inflammation contexts, mainly depending on the cytokine environment. Considering its dual implication, to understand the interactions between the gut microbiota (especially the pathobionts) and IL-22-producing cells, could identify promising targets for IBD therapy.

Here, we identify *Sutterella* sp., as a new pathobiont belonging to the betaproteobacteria class and *the Burkholderiales* order. This anaerobic Gram-negative bacterium can inhibit IL-22 *in vitro* and *in vivo* by regulating AhR activity, without affecting ROR-γt expression. The effectors of activity are secreted by *Sutterella* and are protein-based (>30 kDa).

## RESULTS

### The *Sutterella* sp. bacteria correlate negatively with gut IL-22- and IL-17-producing cells

The gut microbiota is essential to maintain gut IL-22 and IL-17A (IL-17)-producing cells^14, 15^. Therefore, as previously reported^16, 17^, we found that vancomycin or a mix of antibiotics decrease the production of these two cytokines by αβ CD4^+^ T cells, γδ T cells, CD4^+^ CD3 ^neg^ or CD4^neg^ CD3 ^neg^ cells (gated as described in Figure S1A) in the small intestine (SI), and colon (Figures 1A-B; Figures S1B, S1C). Only colonic γδ T cells are differentially regulated since their production of IL-17 and IL-22 were increased with antibiotics treatment (Figures S1B-S1C), as we previously observed^17^.

**Figure 1.**
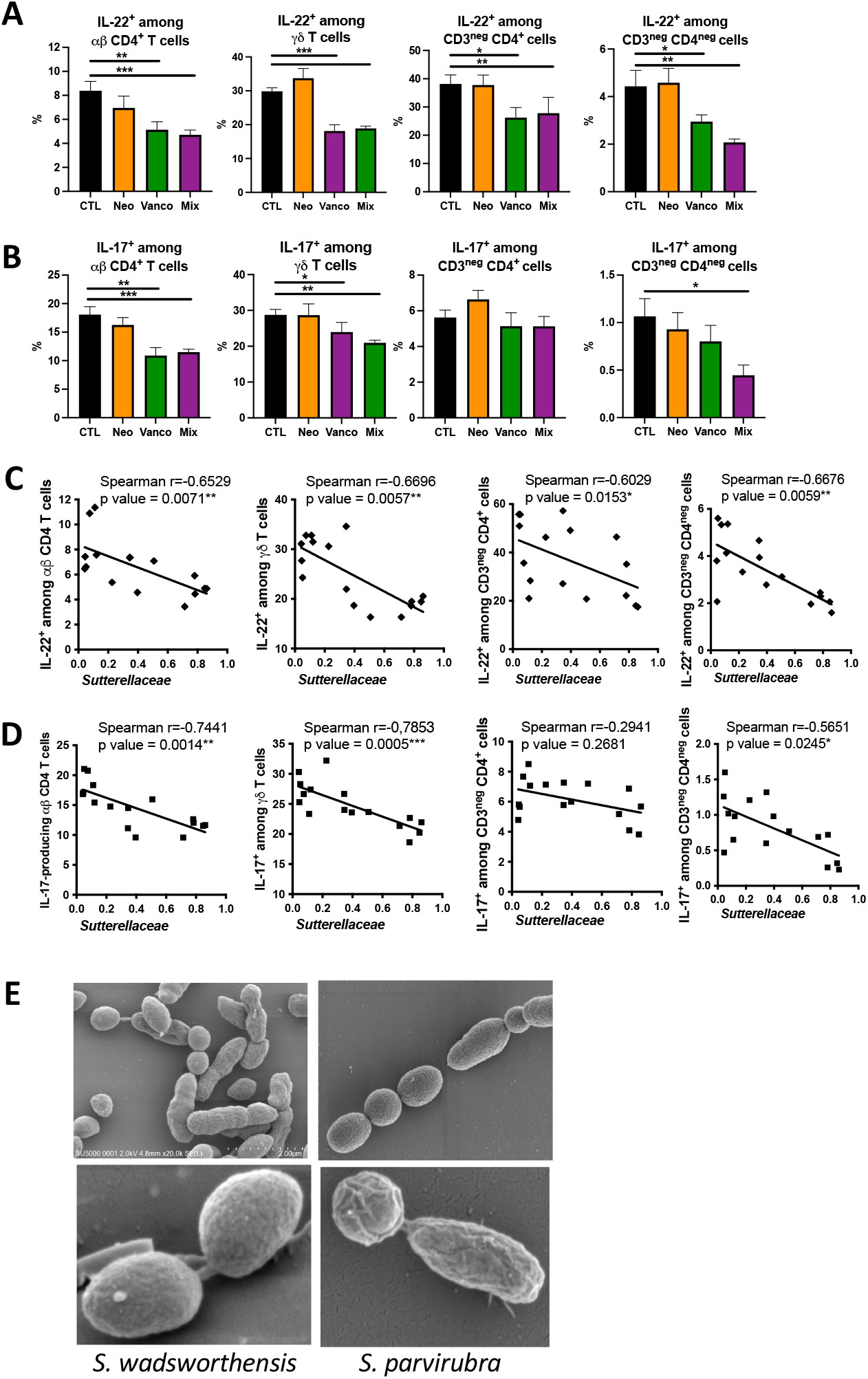
IL-22 and IL-17-producing cells reduce with antibiotics while bacteria from Sutterellaceae family increase. **(A, B)** Percentage of IL-22 (A) and IL-17 (B) among gated CD4^+^ αβ T cells, γδ T cells, CD3_neg_ CD4_+_ and CD3_neg_ CD4_neg_ cells from small intestine obtained from untreated mice (control, CTL), neomycin (Neo)-, vancomycin (Vanco)– or neomycin+vancomycin (Mix)-treated mice. Cells were stimulated with PMA + ionomycin + IL-1β + IL-23 for 4 hours. **(C, D)** Correlation by Spearman r between percentage of IL-22 (C) and IL-17 (D) -producing cells gated as in A and *Sutterellacae* abondance in small intestine. **(E)** SEM images of *S. wadsworthensis* and *S. parvirubra*. Error bars are SEM from 5 to 10 mice per group (A, B) and from 16 mice (C, D). Significant differences were determined using Kruskal-Wallis test (A, B): *p < 0.05; **p < 0.01; ***p < 0.001. See also Figure S1.

Although several commensals and microbiota-derived metabolites have been shown to regulate IL-22- and IL-17-producing cells^12, 14, 17, 18^, we analysed a 16S sequencing in order to identify new candidate commensals species controlling IL-22 and/or IL-17 production in the small intestine and colon of antibiotics-treated and untreated mice. The proportion of IL-17^+^ and IL-22^+^ cells correlated positively with the abundance of Segmented Filamentous Bacteria (SFB, *Candidatus arthromitus*) in the small intestine (Figure S1D), which is in line with the well described stimulatory effect of SFB on Th17 cells in the mouse small intestine, thus validating our approach. Interestingly, we also observed a negative correlation between IL-22 and IL-17 productions and bacteria from the family *Sutterellaceae* (Figures 1C-D; Figures S1E-S1F), suggesting that members from this family could inhibit IL-22 and IL-17 production. Highly prevalent in the human intestinal mucosa^19, 20^, the genus *Sutterella is* increased in the faecal microbiota of patients with UC, and is thus a genus of interest in intestinal inflammation contexts. The two main species of *Sutterella* found in humans are **S. wadsworthensis** and **S. parvirubra**, with a mix of two different morphologies in pure culture: rod and oval shaped (Figure 1E).

### The *Sutterella* sp. bacteria inhibit IL-22 and IL-17 production *in vitro*

To test the effect of *Sutterella* sp. on IL-22 and IL-17 production, we used the culture supernatant of **S. wadsworthensis** and **S. parvirubra** to stimulate the immune cells isolated from the SI lamina propria. **S. wadsworthensis** and **S. parvirubra** supernatants inhibited IL-22 production (Figure 2A) and, to a lower extent, IL-17 production (Figure 2B) by murine immune cells *in vitro*. Flow cytometry analysis revealed that **S. wadsworthensis** inhibited IL-22 and IL-17 production by αβ CD4^+^ T cells, γδ T cells, CD4^+^ CD3 ^neg^ or CD4^neg^ CD3 ^neg^ cells from SI, as early as 4h (Figure 2C). To evaluate the generality of these observations, we analyzed cells from peripheral lymph nodes and found that the proportion and the quantity of IL-22 and IL-17 produced by peripheral γδ T cells and CD4^+^ CD3^neg^ cells, were also inhibited by **S. wadsworthensis** supernatants (Figures 2D-E), without inducing any toxicity (Figure S2A). Within four hours of incubation, **S. parvirubra** only decreased the proportion of IL-22^+^ γδ T cells (Figure S2B) without any toxic effects (Figure S2C). Of note, other members of the Pseudomonadota phylum, such as *E. coli*, or Burkholderiales order, such as *Achromobacter denitrificans*, did not exhibit any inhibitory effect on IL-22 and IL-17 productions (Figure 2F).

**Figure 2.**
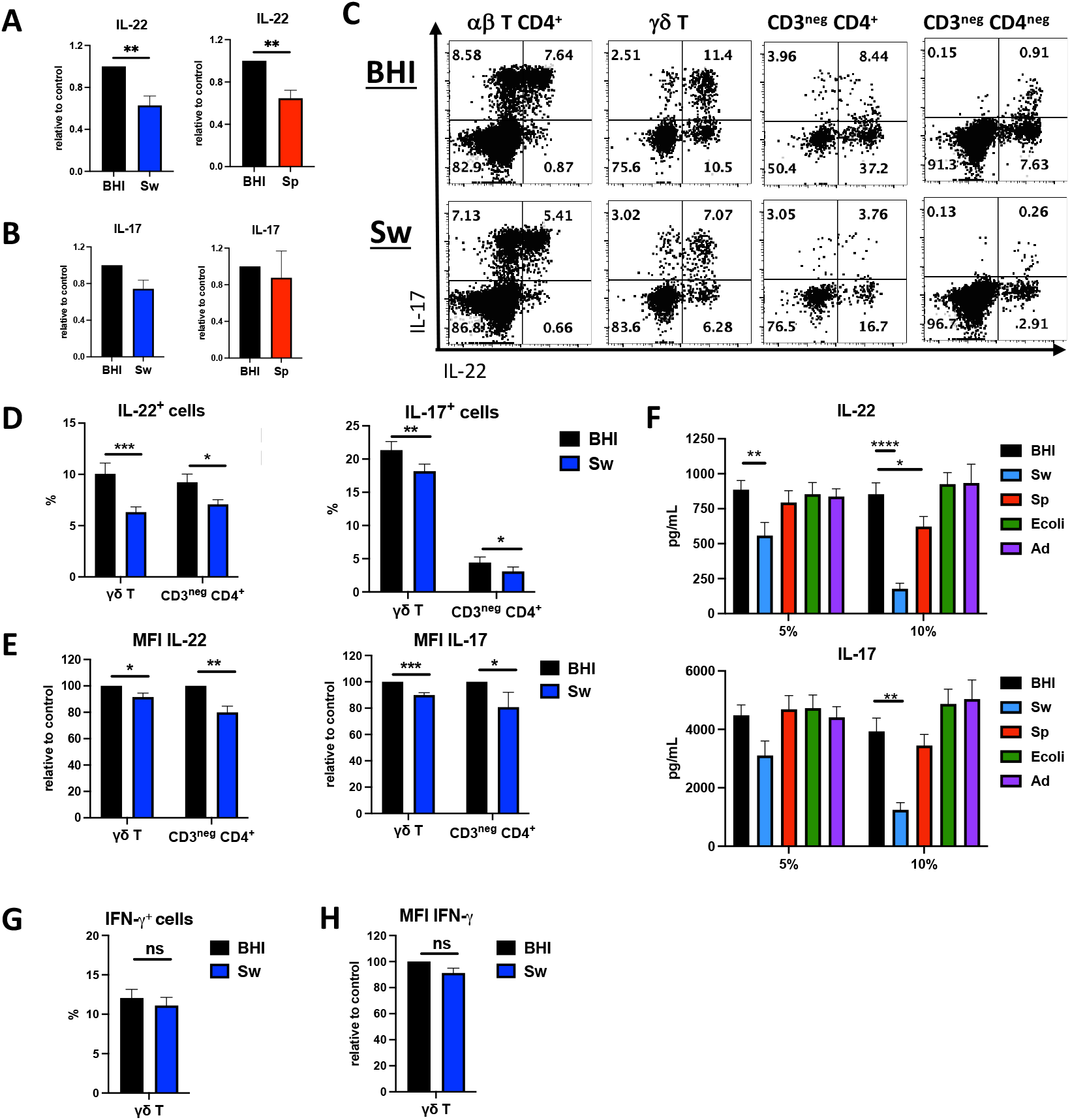
Sutterella regulates IL-17 and IL-22 productions in vitro. **(A, B)** Cells from LP of small intestine were cultured for 24h with BHI (black) or *S. wadsworthensis* (Sw, blue) or *S. parvirubra* (Sp, red) culture supernatant and stimulated with PMA + Ionomycin + IL-1β + IL-23. IL-22 and IL-17 productions were measured in the supernatants. **(C)** Intracellular analysis of IL-17 and IL-22 expression by gated αβ CD4+ T cells, γδ T cells, CD3^neg^ CD4^+^ cells and CD3^neg^ CD4^neg^ cells from small intestine cultured with BHI (top), Sw (bottom) culture supernatant and stimulated for the 3 last hours with PMA + Ionomycin + IL-1β + IL-23. Data are representative of three independent experiments **(D)** Intracellular analysis of IL-22 (left) and IL-17 (right) expression by gated γδ T cells and CD3^neg^ CD4^+^ cells from pLN cultured with BHI (black), Sw (blue) culture supernatant and stimulated as in C. **(E)** Normalized geometric mean fluorescence intensity for IL-22 (left) and IL-17 (right) in gated γδ T cells and CD3^neg^ CD4_+_ cells obtained as described in D. **(F)** Cells from pLN were cultured for 24h with BHI (black), *S. wadsworthensis* (Sw, blue), *S. parvirubra* (Sp, red), *E. coli* (Ecoli, green) or *A. denitrificans* (Ad, violet) culture supernatant and stimulated as in A. IL-22 (top) and IL-17 (bottom) productions were measured in the supernatants. **(G)** Intracellular analysis of IFN-γ expression by gated γδ T cells obtained as described in D. **(H)** Normalized geometric mean fluorescence intensity for IFN-γ in gated γδ T cells obtained as described in G. Error bars are SEM with 8-9 mice (A), 5 mice (B), 13 mice (D-F). ns, not significant, *P < 0.05, **P < 0.005, ***P < 0.001, ****P < 0.0001; by Wilcoxon test or Friedman test. See also Figure S2

Although it has been previously speculated that *Sutterella* sp. has a proinflammatory profile^20^, it does not seem to be the case in our hands. Besides the inhibition of IL-17 and IL-22, **S. wadsworthensis** and **S. parvirubra** did not affect either the production of the pro-Th1 cytokine IFN-γ by peripheral γδ T (Figures 2G-H; Figure S2D), or the release of IL-6, TNF-α and IFN-γ by SI lamina propria cells (Figure S2E). Finally, *Sutterella* did not alter IL-8 production by epithelial cell line, unstimulated or stimulated with TNF-α, in comparison to control (Figure S2F).

### The *Sutterella* sp. bacteria inhibit IL-22 production *in vivo*

In line with *in vitro* results, intragastric gavage *of *S. wadsworthensis** induced a significant reduction in the frequency of IL-22-producing CD4^+^ CD3^neg^ cells in SI (Figure S3A) and decreased production of IL-22 by all populations (αβ CD4^+^ T cells, γδ T cells, CD3^neg^ CD4^+^ and CD3^neg^ CD4^neg^) as demonstrated by geometric mean fluorescence intensity (MFI) analysis (Figure S3B). Regarding IL-17, only its quantity produced by αβ CD4^+^ T cells was reduced in **S. wadsworthensis**-treated compared to control mice (Figure S3C, S3D).

Because dead bacteria pellet has no effect *in vitro* on IL-17 and IL-22 production (Figure S3E), we next analyzed the capacity of *Sutterella* culture supernatants to regulate these cytokines *in vivo*. While the proportion of IL-22-producing αβ CD4^+^ T cells from distal SI and IL-22-producing CD3^neg^ CD4^+^ cells from proximal SI were reduced by **S. wadsworthensis** and **S. parvirubra** culture supernatant (Figure 3A-B, Figure S3F), the quantity of IL-22 produced was reduced for all populations (αβ CD4^+^ T cells, γδ T cells, CD3^neg^ CD4^+^ and CD3^neg^ CD4^neg^) from distal and proximal SI, in both strains-treated mice in comparison to control mice (Figure 3A, 3C; Figure S3G). Thus, *Sutterella* sp. is able to inhibit IL-22 production *in vitro* and *in vivo*. Concerning IL-17, as observed with alive bacteria (Figure S3C), the culture supernatants did not affect the proportion (Figure 3D, Figure S3H) but reduced the quantity of cytokine produced by αβ CD4^+^ T cells from distal SI of **S. wadsworthensis** and **S. parvirubra**-treated mice (Figure 3E, Figure S3I).

**Figure 3.**
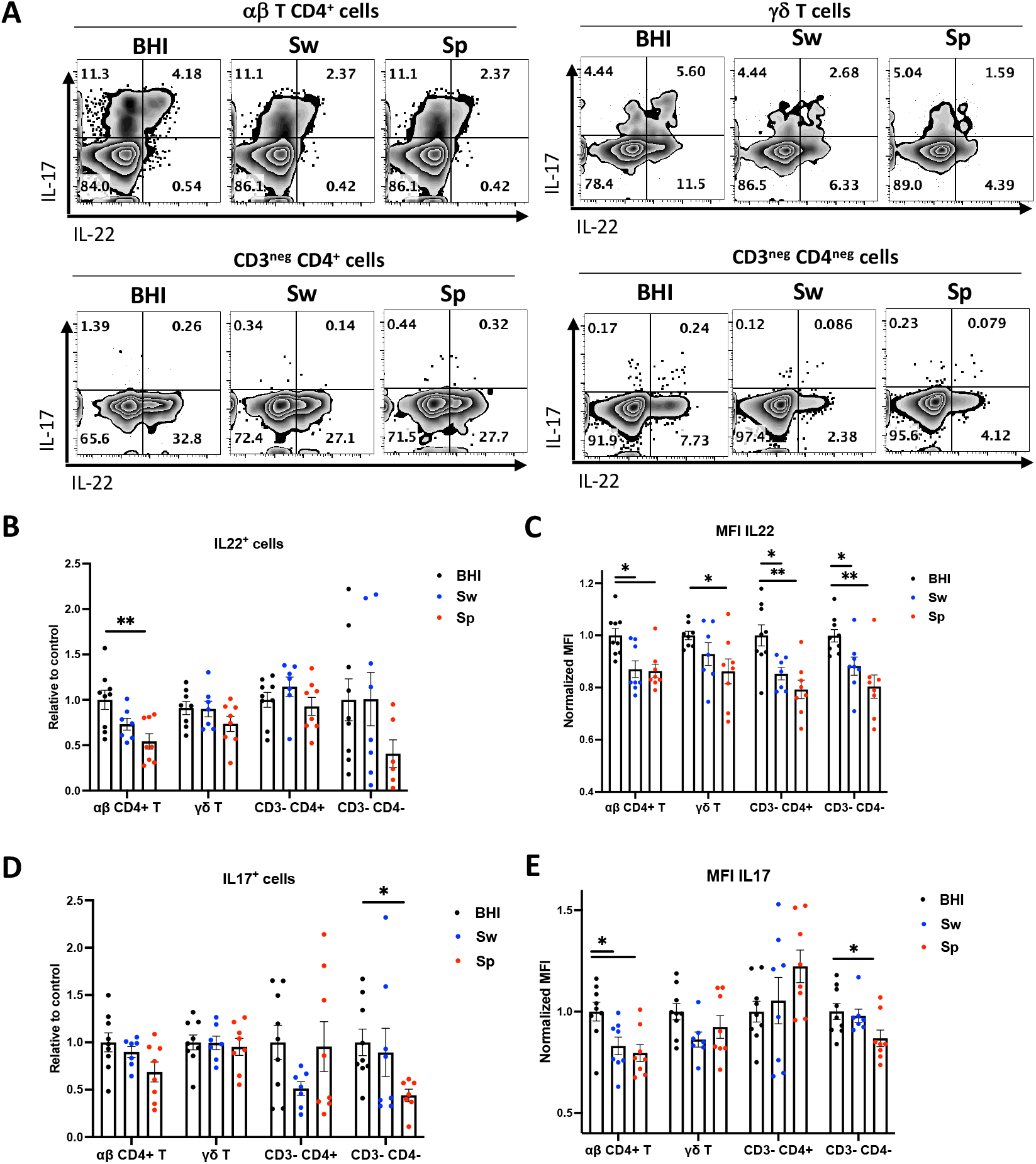
Sutterella regulates IL-17 and IL-22 productions in vivo. **(A)** Representative plots of gated αβ CD4+ T cells (top, left), γδ T cells (top, right), CD3^neg^ CD4+ (bottom left), CD3^neg^ CD4^neg^ (bottom right) cells from distal small intestine obtained from BHI (left), *S. wadsworthensis* (Sw, middle) or *S. parvirubra* (Sp, right) culture supernatant - treated mice. **(B-E)** Intracellular analysis of IL-22 (B) and IL-17 (D) expression and normalized geometric mean fluorescence intensity for IL-22 (C) and IL17 (E) by gated αβ CD4+ T cells, γδ T cells, CD3^neg^ CD4^+^, CD3^neg^ CD4^neg^ cells obtained as in A. In each case, cells were stimulated 3 hours with PMA + Ionomycin + IL-1β + IL-23. Error bars are SEM with 7-9 mice per group for each experiment. *P < 0.05, **P < 0.005, by Kruskal-Wallis test. See also Figure S3

### *Sutterella* sp. bacteria inhibit IL-22 independently of RORγt and STAT3 and likely through AhR

As ROR-γt is required to stimulate IL-22 and IL-17 production^21^, the expression of this transcription factor was assessed after stimulation of cells from pLN with **S. wadsworthensis** and **S. parvirubra** culture supernatant. Although we observed an inhibition of IL-22 mRNA expression following incubation with *Sutterella* supernatants, the expression of RORc gene was maintained (Figure 4A). The proportion of total ROR-γt^+^ cells was not altered by **S. wadsworthensis** and **S. parvirubra** culture supernatant (Figure 4B), as also observed at the γδ T lymphocyte level (Figure S4A). Moreover, among RORγt^+^ ILC3 (CD3^neg^ CD4^+^ RORγt^+^ cells), **S. wadsworthensis** and **S. parvirubra** culture supernatants reduced the proportion of IL-22-competent cells (Figure 4C). Therefore, *Sutterella* sp. acts on IL-22 and IL-17 production independently of RORγt inhibition.

**Figure 4.**
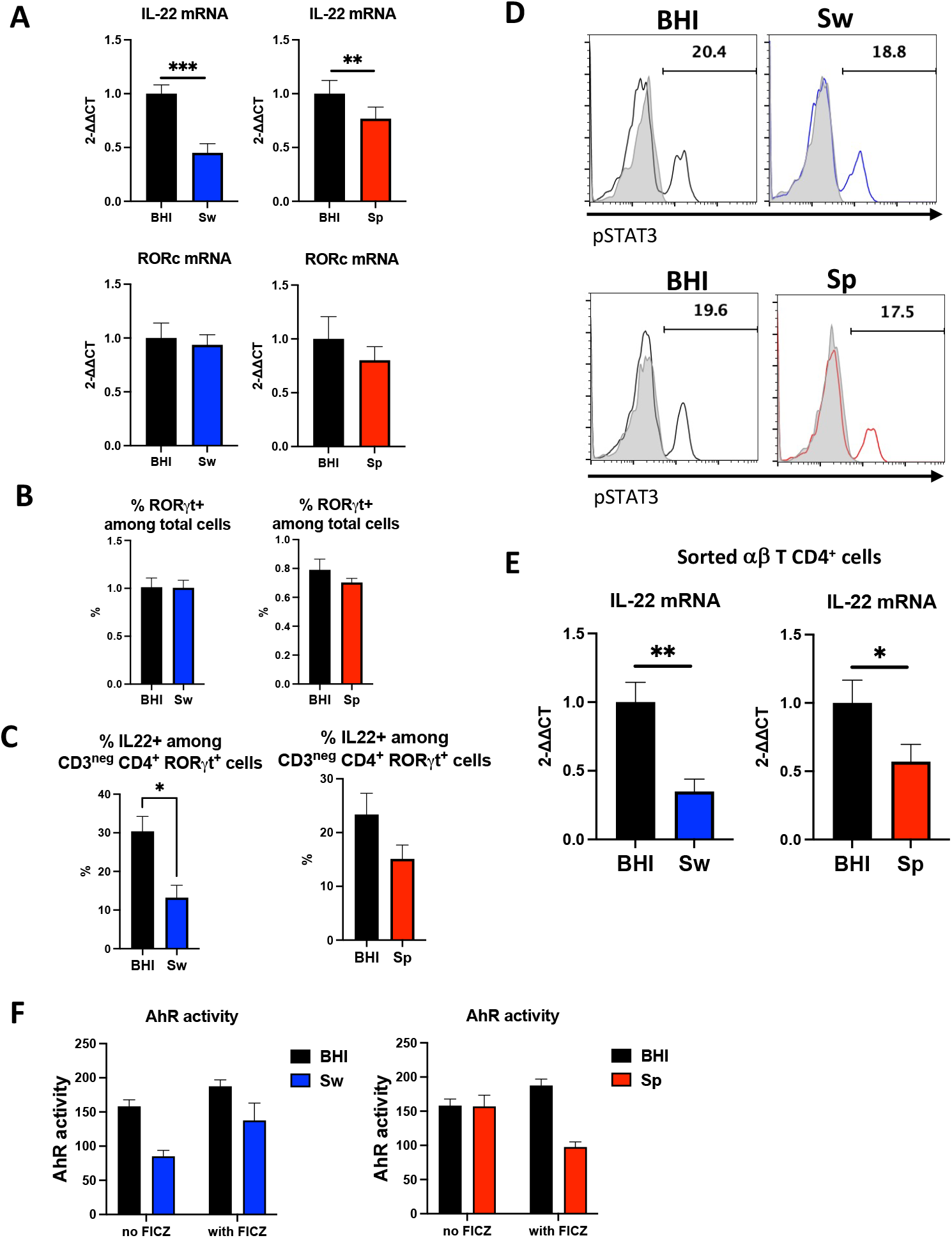
Sutterella regulates directly IL-22 producing cells by the inhibition of AhR. **(A)** Cells from pLN were cultured for 4 hours with BHI (black) or *S. wadsworthensis* (Sw, blue) or *S. parvirubra* (Sp, red) culture supernatant and stimulated with PMA + Ionomycin + IL-1β + IL-23 for the 3 last hours. IL-22 (top) and RORc (bottom) mRNA were quantified by qPCR. **(B)** Proportion of ROR-γt^+^ cells among total cells from LN cells cultured and stimulated as in A. **(C)** Proportion of IL-22^+^ cells among ROR-γt^+^ CD3^neg^ CD4^+^ cells from LN cells cultured and stimulated as in A. **(D)** Flow cytometric detection of intracellular pSTAT3 in gated γδ T cells from pLN, cultured 2h with BHI or *S. wadsworthensis* (Sw, top) or *S. parvirubra* (Sp, bottom) culture supernatant and stimulated with IL-23 for the last hour. Open and shaded areas indicate IL-23 treatment and controls, respectively. **(E)** IL-22 mRNA quantification by qPCR in sorted αβ CD4^+^ T cells cultured 4h with BHI (black) or *S. wadsworthensis* (Sw, blue) or *S. parvirubra* (Sp, red) culture supernatant and stimulated as in A. Mice were treated with DSS and αβ CD4^+^ T cells were sorted from mesenteric LN. **(F)** AhR reporter cell line activation without or with FICZ, in presence of BHI (black) or *S. wadsworthensis* (Sw, blue) or *S. parvirubra* (Sp, red) culture supernatant. Error bars are SEM with 10 mice (A), 6 mice (B, C), 8 mice (D), 7-10 mice (E) and 3 biological replicates (F). *P < 0.05, **P < 0.005, ***P < 0.0005; by Wilcoxon test. See also Figure S4

IL-23 signals are primarily transduced by STAT3^22^. However, IL-23-dependent STAT3 phosphorylation was comparable among cells cultured with the control medium or the supernatants of **S. wadsworthensis** and **S. parvirubra** (Figure 4D and Figure S4B).

Sorted αβ CD4^+^ T cells were examined to assess the direct effect of *Sutterella* supernatants. Upon stimulation, the induced production of IL-22 and IL-17 was inhibited by **S. wadsworthensis** and **S. parvirubra** culture supernatants (Figure 4E and Figure S4C). Thus, *Sutterella* effectively inhibits αβ CD4^+^ T cells functions through direct effects.

Another mechanism that could explain Il-22 and IL-17 inhibition is the modulation of AhR (Aryl hydrocarbon Receptor) pathway^5^. Using an AhR reporter system, we found that **S. wadsworthensis** and **S. parvirubra** supernatants reduced AhR activation (Figure 4F).

### A 30>kDa protein produced by *Sutterella* sp. inhibits IL-22

The *Sutterella* effectors responsible for IL-22 and IL-17 inhibition are secreted molecules, since the culture supernatant has a repressor effect (Figure 2), unlike the bacterial wall (Figure S3E). To assess the nature of these effectors, the supernatants were heated, treated with proteinase K and/or filtered with different molecular weight cutoff membranes. **S. wadsworthensis** culture supernatants heated and treated with proteinase K lost their inhibitory effect on IL-22 production (Figure S5A). Moreover, the fraction below 3 kDa of **S. wadsworthensis** culture supernatant had no effect on IL-22 production by γδ T cells, CD3^neg^ CD4^+^ cells, while the fraction above 3 kDa could inhibit this cytokine (Figure 5A), without affecting cell viability (Figure S5B). The > 30 kDa fraction of **S. wadsworthensis** culture supernatant also significantly reduced IL-22 production (Figure 5B), while the > 30 kDa fractions treated with proteinase and/or heated lost their inhibitory effect on γδ T cells (Figure 5C) and CD3^neg^ CD4^+^ cells (Figure 5D). Such as non-treated culture supernatant, the > 30 kDa fraction had no effect on cell viability (Figure S5C) or IFN-γ production (Figure S5D). Taken together, these results identified that the effectors of **S. wadsworthensis** activity are not metabolites (previously identified in other bacterial commensals)^12, 14, 17, 18^ but proteins larger than 30 kDa.

**Figure 5.**
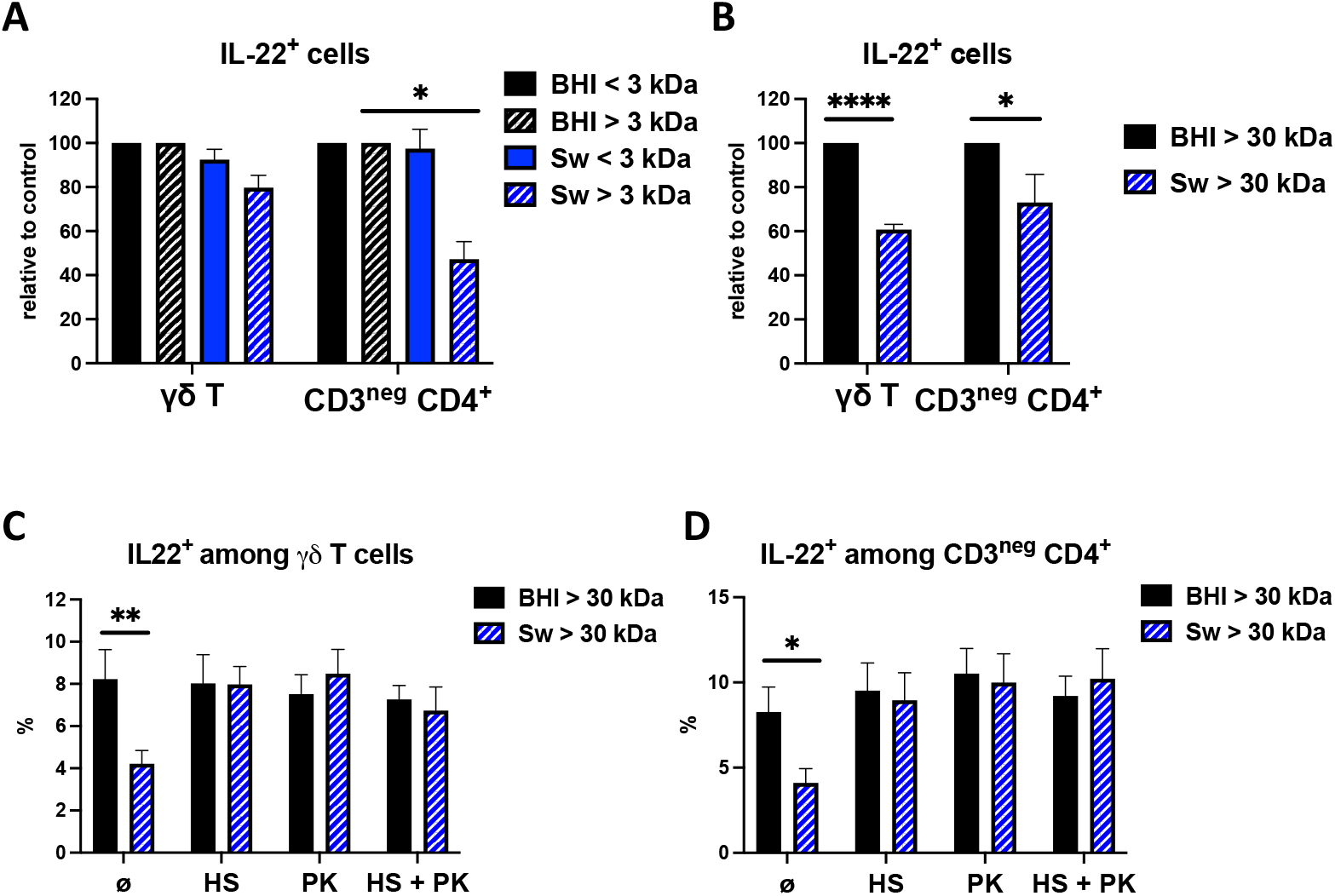
A >30kDa protein from Sutterella regulates IL-22 productions. **(A, B)** Intracellular analysis of IL-22 expression by gated γδ T cells and CD3^neg^ CD4^+^ cells from pLN cultured 4 hours with <3 kDa and >3 kDa fractions (A) or >30 kDa (B) of BHI or Sw culture supernatant and stimulated with PMA + Ionomycin + IL-1β + IL-23 for the 3 last hours. **(C, D)** Intracellular analysis of IL-22 expression by gated γδ T cells (C) and CD3^neg^ CD4_+_cells (D) from pLN cultured 4 hours with >30 kDa of BHI or Sw culture supernatant (pretreated with proteinase K (PK) and/or pre-heated (Heat shock, HS)) and stimulated as in A. Error bars are SEM from 6 mice (A), 7 mice (B) and 5 mice (C, D). Significant differences were determined using paired t test (A) and Wilcoxon test (B-D): *P < 0.05, **P < 0.005, ****P < 0.0001. See also Figure S5

### *Sutterella* sp. inhibits human IL-22 and IL-17

Next, we examined the significance of these results in a human context. We treated peripheral blood mononuclear cells (PBMC) 24h and 48h with **S. wadsworthensis** and **S. parvirubra** supernatants and measured the production of IL-22 and IL-17. As in mice, *Sutterella* inhibit the production of human IL-22 and IL-17 (Figure 6).

**Figure 6.**
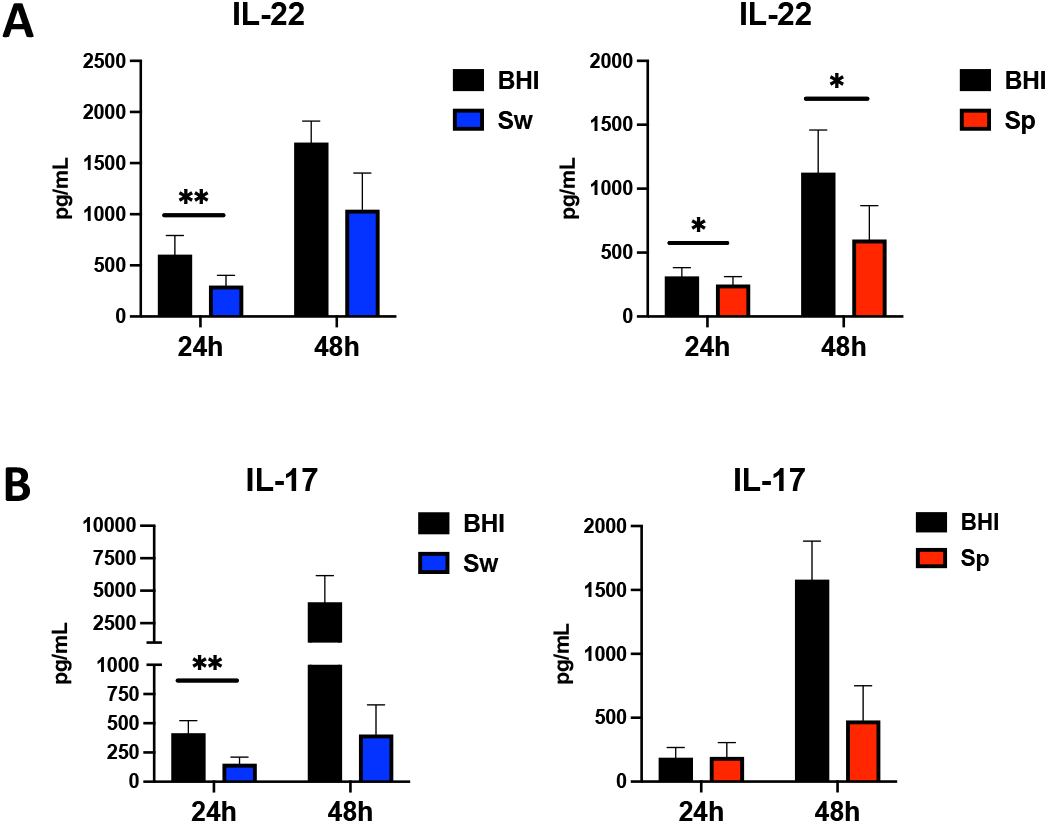
Sutterella regulates IL-17 and IL-22 productions in human. Cells from peripheral blood mononuclear cells (PBMCs) were cultured for 24h or 48h with BHI (black), *S. wadsworthensis* (Sw, blue) or *S. parvirubra* (Sp, red) culture supernatant and stimulated with anti-CD3 and anti-CD28. IL-22 (A) and IL-17 (B) productions were measured in the supernatants.Error bars are SEM from n = 4-9 donors. Significant differences were determined using Wilcoxon test, *P < 0.05, **P < 0.005.

## DISCUSSION

In this study, we observed that antibiotics treatment increases the proportion of betaproteobacteria from *Sutterellaceae* family, which correlates negatively with the proportion of IL-22 and IL-17 producing cells in murine small intestine and colon. *In vitro* and *in vivo* analysis then showed that two species from this bacterial family, **S. wadsworthensis** and **S. parvirubra**, are capable of inhibiting these two cytokines produced by αβ CD4^+^ T cells, γδ T cells, CD4^+^ CD3 ^neg^ or CD4^neg^ CD3 ^neg^ cells. These findings are relevant in human since *Sutterella* sp. also reduces the level of IL-22 and IL-17 secreted by human PBMCs.

The regulation of IL-22 production is primarily associated to gut microbiota-derived metabolites, such as tryptophan metabolites^12^, short-chain fatty acids^17, 18^ and secondary bile acids^23^. Here, we demonstrated that the bio-active effectors of *Sutterella* sp. are > 30 kDa proteins that act directly on lymphoid cells. The production of IL-22 can be regulated through several signaling pathways. IL-23 signal is primarily transduced by STAT3^22^. However, IL-23-dependent STAT3 phosphorylation was comparable between cells incubated with *Sutterella* culture supernatant and culture medium. AhR is another major transcription factor, essential for IL-22 production by ILC3^24^, Th17 cells^25^ and γδ T cells^26^. It can modulate Th17 development independently to ROR-γt expression and STAT3 phosphorylation^27^. However, AhR does not act alone but cooperates with other transcription factors to induce IL-17 or IL-22 production. AhR facilitates notably ROR-γt recruitment at the IL-22 locus to induce its expression^28^. In line with this, our data suggest that *Sutterella* sp. inhibits IL-22 and IL-17 productions, by reducing AhR activity without affecting ROR-γt expression.

Although it has been shown that **S. wadsworthensis** has a mild pro-inflammatory activity, we did not observe the induction of IL-8 by epithelial cells or pro-Th1 cytokines by immune cells in presence of *Sutterella* culture supernatant. Therefore, instead of directly triggering inflammation, *Sutterella* impairs the intestinal immune response, notably by inhibiting IL-22 and IL-17 productions, and could lead to a favorable environment for the invasion of epithelial cells by other pathobionts. In line with this, Moon et al. showed in 2015 the ability of *Sutterella sp*. to degrade IgA antibodies, thus disrupting the mucosal defense^29^.

Patients with UC exhibit an increased amount of *Sutterella* in their fecal microbiota^30^ and a reduced number of IL-22-producing cells in actively inflamed tissues^31^, suggesting a link between *Sutterella* and IL-22 in humans and in IBD. Given the beneficial role of IL-22 in IBD^4^, *Sutterella* sp. may have a deleterious effect and represents a promising target for IBD therapy. This is also a potential biomarker for IBDs diagnosis. In exploring this promising hypothesis and identifying the implicated bio-active proteins, we will pave the way to develop innovative clinical strategies to control gut inflammation.

## Acknowledgments

We thank the employees of the animal facility at the IERP platform, INRAE, UE0907 (Jouy-en-Josas, France) for their technical assistance. We also acknowledge Marie-Laure Aknin, CYM core facility from the “Unité Mixte de Service – Ingénierie et Plateformes au Service de l’Innovation” (UMS-IPSIT, Paris Saclay, France), for access to its equipment and infrastructures and technical assistance. We thank Ambrine Farfar, Rand Fatouh, Chloé Michaudel, Julien Planchais, Ugo Sardo, Mathias L. Richard and Vlad Costache (MIMA2 core facility, INRAE, Jouy-en-Josas, France) for assistance. L.D. was supported by the Foundation for Medical Research.

## Author contributions

L.D., H.S. and M.-L.M. designed the study, analyzed and interpreted the data. L.D., G.O., R.G., A.B., M.B. and M.-L.M. performed experiments. G.D.C., A.M. and L.C. provided technical help. L.D., R.G., A.B., M.B., N.R., C.D., Z.D., P.S., H.S. and M.-L.M. discussed the experiments and results. H.S. and M.-L.M. wrote the manuscript. All authors read and approved the final manuscript.

## Declaration of Interests

The authors declare no competing interests.

## MATERIELS AND METHODS

### Mice

C57Bl/6J mice (7-9-week-old females) were bred and maintained under specific pathogen-free conditions. Animal experiments were performed according to local ethical panel and the Ministère de l’Education Nationale, de l’Enseignement Supérieur et de la Recherche, France under agreement Apafis#20977-2019041215549049.

## Human samples

PBMCs were isolated from the peripheral blood of healthy donors obtained from Etablissement Français du Sang (EFS).

### Bacterial culture

*Escherichia coli* (K-12/MG1655), *Sutterella wadsworthensis* (DSM 14016), isolated from human abdominal fluid, and *Sutterella parvirubra* (DSM 19354), isolated from human faeces, were grown at 37°C in LYHBHI medium [Brain–heart infusion medium supplemented with 0.5% yeast extract (Difco) and 5 mg/liter hemin] supplemented with Formate (1.8 mg/ml; Sigma), Fumarate (1.8 mg/ml; Sigma), cysteine (0.5 mg/ml; Sigma), vitamin K1 (0.0001%, Sigma) and vitamin K3 (3.2 µg/ml, Sigma) in an anaerobic chamber. *Achromobacter denitrificans* (DSM 30026), isolated from soil, was grown at 37°C in the same medium, with agitation in aerobic conditions.

Strains are cultured for 18h, until the end of exponential growth phase/beginning of stationary growth phase. Cell concentration is measured with a flow cytometer (CytoFLEX, Beckman Coulter). After a centrifugation for 15 minutes at 4500 rpm, 4°C, in flying buckets, the supernatants are filtered in 0.2 µm membrane and aliquoted for storage at -20°C. Bacterial cell pellets are resuspended in PBS at a concentration of 1×10^9^ cells/ml, then heat killed at 70°C for 30 minutes. These suspensions are aliquoted for storage at -20°C.

### Antibiotic and *Sutterella* treatments

Mice were treated with vancomycin (500 mg/L; Mylan) and/or neomycin (1 mg/mL; Euromedex) in the drinking water and water solutions were prepared and changed every three days. Fluid intake was monitored and the antibiotic solution was changed every 3 days. In some experiments, mice were pre-treated 4 days with streptomycin sulfate salt (5mg/mL, Sigma) in the drinking water and then inoculated daily via intragastric gavage with a bacterial suspension of 10^9^ CFU/mL in 200 μL of PBS/glycerol or control medium (PBS/glycerol) for 3 weeks. In some other experiments, mice were inoculated daily via oral gavage with 200 μL of filtrated bacterial supernatant or control medium, during 1 week.

### DSS treatment

To obtain IL-22-producing CD4^+^ αβ T cells from mesenteric lymph nodes, mice were administrated drinking water supplemented with 2% DSS (MP Biomedicals) for 7 days, and then allowed to recover by drinking supplemented water for the next 2 days.

### Fractionation or treatment of Sutterella supernatants

Heat shock treatment at 90°C for 35 minutes was performed. **S. wadsworthensis*, *S. parvirubra** culture supernatants and LYHBHI medium were also treated with proteinase K (Qiagen) overnight at 56°C, and finally heated for 20 minutes at 96°C to inactivate the enzyme. In some experiments **S. wadsworthensis*, *S. parvirubra** culture supernatants and LYHBHI medium were fractionated by size using 3-kDa-molecular-weight-cutoff (MWCO) or 30-kDa-MWCO centrifugal filters (Amicon). Filters were rinsed once with PBS 1% BSA (Sigma). Then, 3 mL of bacterial culture supernatants or LYHBHI medium were added to the filter and centrifuged at 3,000 g at 4°C. The flowthroughs and the top fractions were resuspended in LYHBHI medium up to 3 mL.

### Fecal and ileal fluid DNA extraction

Fecal and ileal fluid DNA were extracted from the weighted samples as previously described^12.^

### Murine cell preparation

Isolation of mononuclear cells from mouse lamina propria: small intestine between stomach and cecum, large intestine between cecum and anal verge were cut out and open longitudinally. Tissues were washed with cold PBS1X to remove the fecal and then were cut cross-sectionally into 0.5–1 cm long pieces and then mixed with 5 mL pre-warmed HBSS/FCS/EDTA/DTT in the shaker at 37°C for 20 min. The supernatants were discarded and pellets were washed with cold PBS1X. The tissue pieces were then transferred to a new tube and digested with collagenase type IV (Gibco) or the mouse Lamina Propria Dissociation Kit (Miltenyi) and DNAse (Roche) for 30 min. All the contents were passed through a 100 μm cell strainer. LPLs were obtained using the 40/80 Percoll centrifugation (GE Healthcare). Mouse lymph nodes preparation: peripheral lymph nodes (pLN, axillary, brachial, and inguinal) or mesenteric lymph nodes (mLN) were homogenized and washed in RPMI 1640 medium (Gibco) + 10% (vol/vol) FCS (Sigma-Aldrich) + 1% Hepes (Gibco) + 100 U/mL Penicillin and 100 μg/mL Streptomycin (Gibco).

### Human PBMCs isolation

Heparinized whole blood was diluted with PBS 1:1, layered over Histopaque-1077 (Sigma-Aldrich) and centrifuged at 400g for 20 min without brake at room temperature. The peripheral blood mononuclear cell (PBMC) fraction was then washed in cold PBS and used for *in vitro* culture assays.

### *In vitro* Culture Assays and stimulation

Murine cells and human PBMCs were cultured in RPMI 1640 medium (Gibco) supplemented with 10% (vol/vol) FCS (Sigma-Aldrich) + 1% Hepes (Gibco) + 100 U/mL Penicillin and 100 μg/ml Streptomycin (Gibco), in the presence of BHI or *Sutterella* supernatant at 2, 5 or 10% or dead *Sutterella* (MOI 10:1), during 4 or 24 hours. Murine cells were prepared at 5×10^5^ cells (at 2.5×10^6^ cells/mL) per 96-well plate (CellStar) and stimulated for the 3 or 23 last hours in culture medium with 50 ng/mL phorbol 12myristate 13-acetate (Sigma-Aldrich), 1 mg/mL ionomycin (Sigma-Aldrich), 20 ng/mL mouse IL-23 recombinant protein and 10 ng/mL mouse IL-1β recombinant protein (R&D Systems). PBMCs were prepared at 1×10^6^ cells (at 1×10^6^ cells/mL) per 24-well plate (CellStar) and stimulated 24h in culture medium with coated anti-human CD3 (2 μg/mL, Biolegend) and soluble anti-human CD28 (4 μg/mL, Biolegend). The culture supernatants were frozen at -20°C until processing.

HT29 cell line (ATCC HTB-38) was cultured in Dulbecco’s Modified Eagle Medium (DMEM, Gibco) with glutamax supplemented with 10% (vol/vol) FCS (Sigma-Aldrich) + 100 U/mL Penicillin and 100 μg/ml Streptomycin (Gibco), at 37°C in a 5% CO2 atmosphere. For some experiments, cell lines (5×10^5^ cells per well, plated 72 hours before the experiment in 24-well plate) were cultured with BHI or *Sutterella* supernatant at 5% and stimulated 24 hours with human recombinant TNF-α (5 ng/mL, Preprotech).

### Flow Cytometry and Cell Sorting

Flow cytometry was carried out by using BD-LSRII machines. αβ CD4^+^ T cells were sorted by using BD-Aria machine or mouse CD4^+^ T Cell Isolation Kit (Miltenyi).

Single cell suspensions were prepared in FACS buffer (PBS + 2% (vol/vol) FCS + 0.01% (vol/vol) sodium azide; Sigma-Aldrich). Cells were stained on ice in PBS 1X (GIBCO) with Fixable Viability Dye eFluor 506 (eBioscience) or Zombie Aqua Fixable Viability Kit (BioLegend). Cells were surface stained in FACS buffer with the following antibodies ; from eBioscience: FITC–labeled anti-CD3ε (145-2C11), PE-labeled anti-CD4 (RM4-5), anti-CD16/32 (93), PECy7-labeled anti–IFN-γ (XMG1.2), eF450-labeled anti-IL-17 (17B7), PECy7-labeled anti-ROR-γT (B20); from BioLegend: APC-labeled anti-TCRγδ (GL3)PE-labeled anti-IL-22 (Poly5164), BV785-labeled anti-CD4 (RM4-5), BV605-labeled anti-CD8α (53-6.7). Cells were washed in FACS buffer before analysis. In mouse, γδ T cells are CD3^+^TCRγδ^+^ CD4^neg^ CD8^neg^, CD4^+^ T lymphocytes are CD3^+^TCRγδ_neg_ CD4^+^ CD8^neg^, CD4^+^ CD3^neg^ cells are γδ TCR^neg^ CD3^neg^ CD4^+^ CD8^neg^ and CD4^neg^ CD3^neg^ cells are γδ TCR^neg^ CD3^neg^ CD4^neg^ CD8^neg^.

For intracellular analysis of IL-17, IL-22 and IFN-γ, mouse cells were first stimulated in the presence of 10 μg/mL Brefeldin-A (Sigma-Aldrich). After surface staining, cells were fixed and permeabilized by using 4% (wt/vol) PFA in conjunction with 0.5% saponin in PBS1X. We performed intracellular staining for ROR-γt according to the manufacturers’ instructions (FOXP3/Transcription Factor Staining Buffer Set, Invitrogen). Data were analysed by using FlowJo software.

### RNA Expression Analysis

Total RNA was isolated with RNeasy Micro Kit or Mini Kit (Qiagen) as per manufacturer’s instructions. For real-time RT-PCR analysis, RNA was reverse transcribed with LunaScript® RT SuperMix kit (New England Biolabs) by a mix of random hexamers and oligos-dT primers. Quantitative PCR was performed with Luna® Universal qPCR Master Mix (NEB) on StepOne equipment (Applied Biosystems). Relative expression is displayed in arbitrary units normalized to GAPDH via ΔΔCt method. Primer sequences are available upon request.

### Cytokine quantification

ELISAs were performed on the supernatants to quantify mouse IL-22 and IL-17A cytokines (Invitrogen) and human IL-8 cytokine (R&D Systems), according to the manufacturer’s instructions, with Tween 20 (Sigma-Aldrich) and H2SO4 (VWR Chemicals). Legendplex (Mouse Th17 Panel, BioLegend) assay on cell supernatants was performed according to the manufacter recommendations.

### AhR activity measurement

The AhR activity was measured using HepG2-Lucia™ AhR reporter cells (InvivoGen, France) as described before^12^. FICZ (6-Formylindolo[3,2-b]carbazole) was purchased from Enzo Life Sciences.

### 16S sequencing

Bacterial DNA was extracted from the weighted stool samples or ileal fluids as previously described^12^. 16S rDNA gene sequencing was performed as previously described^12^.

### Scanning electron microscopy

50 µL of the bacterial culture were directly fixed in 2 mL volume of 2 % glutaraldehyde buffered with sodium cacodylate 0.1M, during 2 h at room temperature then overnight at 4°C, in a 24-well plate (TPP 92024, Switzerland) that contains 1cm x 1cm glass slides at the bottom of the well. The glass slides were previously cleaned by 70% Ethanol, 10 min plasma cleaner, then cut with a diamond pen, sonicated 15min in 100% Ethanol, then in MiliQ water, and coated with 0,01% Poly-L-lysin (Sigma Merck P4707-50mL) during 5 min and finally rinsed with MiliQ water. Samples thus attached to the glass slides at the bottom of the wells were rinsed twice during 10 min in 0.2 M sodium cacodylate buffer, then in successive baths of ethanol (50, 70, 90, 100, and anhydrous 100 %), and finally dried using a Leica EM300 critical point apparatus with slow 20 exchange cycles, and 2 min delay between steps. Samples were mounted on aluminum stubs with adhesive carbon (EMS, LFG France) and coated with 6 nm of Au/Pd using a Quorum SC7620, 50 Pa of Ar, 180 s of sputtering at 3.5 mA. Samples were observed using the SE detector of a FEG SEM Hitachi SU5000, 2 KeV, 30 spot size, 5 mm working distance. All the preparation steps were done using the MIMA2 core facility, INRAE, Jouy-en-Josas, France (https://doi.org/10.15454/1.5572348210007727E12).

### Quantification and statistical analysis

Results are expressed as the mean ± standard error of the mean (SEM). All statistical analysis was performed using GraphPad Prism 10 software, using two-tailed Man-Whitney test or Wilcoxon test to compare two groups; and ordinary one-way ANOVA or Friedman test to compare 3 groups. Correlation significance was determined using linear regression. *, p < 0.05; **, p < 0.01; ***, p < 0.001. n represents the number of mice or human donors per group. Statistical details of experiments and exact value of n can be found in the figure legends.

## Notes

### Competing Interest Statement

The authors have declared no competing interest.

## REFERENCES

1. Zheng D, Liwinski T, Elinav E. Interaction between microbiota and immunity in health and disease. Cell Res. 2020 Jun;30(6):492–506. doi: 10.1038/s41422-020-0332-7.

2. Yue B, Yu ZL, Lv C, Geng XL, Wang ZT, Dou W. Regulation of the intestinal microbiota: An emerging therapeutic strategy for inflammatory bowel disease. World J Gastroenterol. 2020 Aug 14;26(30):4378–4393. doi: 10.3748/wjg.v26.i30.4378.

3. Gilliland A, Chan JJ, De Wolfe TJ, Yang H, Vallance BA. Pathobionts in Inflammatory Bowel Disease: Origins, Underlying Mechanisms, and Implications for Clinical Care. Gastroenterology. 2024 Jan;166(1):44–58. doi: 10.1053/j.gastro.2023.09.019

4. Mizoguchi A, Yano A, Himuro H, Ezaki Y, Sadanaga T, Mizoguchi E. Clinical importance of IL-22 cascade in IBD. J Gastroenterol. 2018 Apr;53(4):465–474. doi: 10.1007/s00535-017-1401-7.

5. Zenewicz LA. IL-22: There Is a Gap in Our Knowledge. Immunohorizons. 2018 Jul 5;2(6):198–207. doi: 10.4049/immunohorizons.1800006

6. Klotskova HB, Kidess E, Nadal AL, Brugman S. The role of interleukin-22 in mammalian intestinal homeostasis: Friend and foe. Immun Inflamm Dis. 2024 Feb;12(2):e1144. doi: 10.1002/iid3.1144.

7. Zheng Y, Valdez PA, Danilenko DM, Hu Y, Sa SM, Gong Q, Abbas AR, Modrusan Z, Ghilardi N, de Sauvage FJ, Ouyang W. Interleukin-22 mediates early host defense against attaching and effacing bacterial pathogens. Nat Med. 2008 Mar;14(3):282–9. doi: 10.1038/nm1720.

8. Hasegawa M, Yada S, Liu MZ, Kamada N, Muñoz-Planillo R, Do N, Núñez G, Inohara N. Interleukin-22 regulates the complement system to promote resistance against pathobionts after pathogen-induced intestinal damage. Immunity. 2014 Oct 16;41(4):620–32. doi: 10.1016/j.immuni.2014.09.010.

9. Behnsen J, Jellbauer S, Wong CP, Edwards RA, George MD, Ouyang W, Raffatellu M. The cytokine IL-22 promotes pathogen colonization by suppressing related commensal bacteria. Immunity. 2014 Feb 20;40(2):262–73. doi: 10.1016/j.immuni.2014.01.003.

10. Muñoz M, Heimesaat MM, Danker K, Struck D, Lohmann U, Plickert R, Bereswill S, Fischer A, Dunay IR, Wolk K, Loddenkemper C, Krell HW, Libert C, Lund LR, Frey O, Hölscher C, Iwakura Y, Ghilardi N, Ouyang W, Kamradt T, Sabat R, Liesenfeld O. Interleukin (IL)-23 mediates Toxoplasma gondii-induced immunopathology in the gut via matrixmetalloproteinase-2 and IL-22 but independent of IL-17. J Exp Med. 2009 Dec 21;206(13):3047–59. doi: 10.1084/jem.20090900.

11. Fang L, Pang Z, Shu W, Wu W, Sun M, Cong Y, Liu Z. Anti-TNF Therapy Induces CD4+ T-Cell Production of IL-22 and Promotes Epithelial Repairs in Patients With Crohn’s Disease. Inflamm Bowel Dis. 2018 Jul 12;24(8):1733–1744. doi: 10.1093/ibd/izy126.

12. Lamas B, Richard ML, Leducq V, Pham HP, Michel ML, Da Costa G, Bridonneau C, Jegou S, Hoffmann TW, Natividad JM, Brot L, Taleb S, Couturier-Maillard A, Nion-Larmurier I, Merabtene F, Seksik P, Bourrier A, Cosnes J, Ryffel B, Beaugerie L, Launay JM, Langella P, Xavier RJ, Sokol H. CARD9 impacts colitis by altering gut microbiota metabolism of tryptophan into aryl hydrocarbon receptor ligands. Nat Med. 2016 Jun;22(6):598–605. doi: 10.1038/nm.4102.

13. Kamanaka M, Huber S, Zenewicz LA, Gagliani N, Rathinam C, O’Connor W Jr, Wan YY, Nakae S, Iwakura Y, Hao L, Flavell RA. Memory/effector (CD45RB(lo)) CD4 T cells are controlled directly by IL-10 and cause IL-22-dependent intestinal pathology. J Exp Med. 2011 May 9;208(5):1027–40. doi: 10.1084/jem.20102149.

14. Gaboriau-Routhiau V, Rakotobe S, L. cuyer E, Mulder I, Lan A, Bridonneau C, Rochet V, Pisi A, De Paepe M, Brandi G, Eberl G, Snel J, Kelly D, Cerf-Bensussan N. The key role of segmented filamentous bacteria in the coordinated maturation of gut helper T cell responses. Immunity. 2009 Oct 16;31(4):677–89. doi: 10.1016/j.immuni.2009.08.020

15. Ekmekciu I, von Klitzing E, Fiebiger U, Escher U, Neumann C, Bacher P, Scheffold A, Kühl AA, Bereswill S, Heimesaat MM. Immune Responses to Broad-Spectrum Antibiotic Treatment and Fecal Microbiota Transplantation in Mice. Front Immunol. 2017 Apr 19;8:397. doi: 10.3389/fimmu.2017.00397

16. Ivanov II, Frutos Rde L, Manel N, Yoshinaga K, Rifkin DB, Sartor RB, Finlay BB, Littman DR. Specific microbiota direct the differentiation of IL-17-producing T-helper cells in the mucosa of the small intestine. Cell Host Microbe. 2008 Oct 16;4(4):337–49. doi: 10.1016/j.chom.2008.09.009.

17. Dupraz L, Magniez A, Rolhion N, Richard ML, Da Costa G, Touch S, Mayeur C, Planchais J, Agus A, Danne C, Michaudel C, Spatz M, Trottein F, Langella P, Sokol H, Michel ML. Gut microbiota-derived short-chain fatty acids regulate IL-17 production by mouse and human intestinal γδ T cells. Cell Rep. 2021 Jul 6;36(1):109332. doi: 10.1016/j.celrep.2021.109332

18. Fachi JL, Sécca C, Rodrigues PB, Mato FCP, Di Luccia B, Felipe JS, Pral LP, Rungue M, Rocha VM, Sato FT, Sampaio U, Clerici MTPS, Rodrigues HG, Câmara NOS, Consonni SR, Vieira AT, Oliveira SC, Mackay CR, Layden BT, Bortoluci KR, Colonna M, Vinolo MAR. Acetate coordinates neutrophil and ILC3 responses against C. difficile through FFAR2. J Exp Med. 2020 Mar 2;217(3):jem.20190489. doi: 10.1084/jem.20190489.

19. Mukhopadhya I, Hansen R, Nicholl CE, Alhaidan YA, Thomson JM, Berry SH, Pattinson C, Stead DA, Russell RK, El-Omar EM, Hold GL. A comprehensive evaluation of colonic mucosal isolates of Sutterella wadsworthensis from inflammatory bowel disease. PLoS One. 2011;6(10):e27076. doi: 10.1371/journal.pone.0027076.

20. Hiippala K, Kainulainen V, Kalliomäki M, Arkkila P, Satokari R. Mucosal Prevalence and Interactions with the Epithelium Indicate Commensalism of Sutterella spp. Front Microbiol. 2016 Oct 26;7:1706. doi: 10.3389/fmicb.2016.01706.

21. Ivanov II, McKenzie BS, Zhou L, Tadokoro CE, Lepelley A, Lafaille JJ, Cua DJ, Littman DR. The orphan nuclear receptor RORgammat directs the differentiation program of proinflammatory IL-17+ T helper cells. Cell. 2006 Sep 22;126(6):1121–33. doi: 10.1016/j.cell.2006.07.035.

22. Parham C, Chirica M, Timans J, Vaisberg E, Travis M, Cheung J, Pflanz S, Zhang R, Singh KP, Vega F, To W, Wagner J, O’Farrell AM, McClanahan T, Zurawski S, Hannum C, Gorman D, Rennick DM, Kastelein RA, de Waal Malefyt R, Moore KW. A receptor for the heterodimeric cytokine IL-23 is composed of IL-12Rbeta1 and a novel cytokine receptor subunit, IL-23R. J Immunol. 2002 Jun 1;168(11):5699–708. doi: 10.4049/jimmunol.168.11.5699.

23. Hang S, Paik D, Yao L, Kim E, Trinath J, Lu J, Ha S, Nelson BN, Kelly SP, Wu L, Zheng Y, Longman RS, Rastinejad F, Devlin AS, Krout MR, Fischbach MA, Littman DR, Huh JR. Bile acid metabolites control TH17 and Treg cell differentiation. Nature. 2019 Dec;576(7785):143–148. doi: 10.1038/s41586-019-1785-z. Epub 2019 Nov 27. Erratum in: Nature. 2020 Mar;579(7798):E7. doi: 10.1038/s41586-020-2030-5.

24. Lee JS, Cella M, McDonald KG, Garlanda C, Kennedy GD, Nukaya M, Mantovani A, Kopan R, Bradfield CA, Newberry RD, Colonna M. AHR drives the development of gut ILC22 cells and postnatal lymphoid tissues via pathways dependent on and independent of Notch. Nat Immunol. 2011 Nov 20;13(2):144–51. doi: 10.1038/ni.2187.

25. Gaffen SL, Jain R, Garg AV, Cua DJ. The IL-23-IL-17 immune axis: from mechanisms to therapeutic testing. Nat Rev Immunol. 2014 Sep;14(9):585–600. doi: 10.1038/nri3707.

26. Martin B, Hirota K, Cua DJ, Stockinger B, Veldhoen M. Interleukin-17-producing gammadelta T cells selectively expand in response to pathogen products and environmental signals. Immunity. 2009 Aug 21;31(2):321–30. doi: 10.1016/j.immuni.2009.06.020.

27. Kimura A, Naka T, Nohara K, Fujii-Kuriyama Y, Kishimoto T. Aryl hydrocarbon receptor regulates Stat1 activation and participates in the development of Th17 cells. Proc Natl Acad Sci U S A. 2008 Jul 15;105(28):9721–6. doi: 10.1073/pnas.0804231105.

28. Gutiérrez-Vázquez C, Quintana FJ. Regulation of the Immune Response by the Aryl Hydrocarbon Receptor. Immunity. 2018 Jan 16;48(1):19–33. doi: 10.1016/j.immuni.2017.12.012.

29. Moon C, Baldridge MT, Wallace MA, D CA, Burnham, Virgin HW, Stappenbeck TS. Vertically transmitted faecal IgA levels determine extra-chromosomal phenotypic variation. Nature. 2015 May 7;521(7550):90–93. doi: 10.1038/nature14139. Epub 2015 Feb 16.

30. Paramsothy, S. et al. Specific Bacteria and Metabolites Associated With Response to Fecal Microbiota Transplantation in Patients With Ulcerative Colitis. Gastroenterology 156, 1440–1454 e1442, doi:10.1053/j.gastro.2018.12.001 (2019).

31. Leung, J. M. et al. IL-22-producing CD4+ cells are depleted in actively inflamed colitis tissue. Mucosal Immunol 7, 124–133, doi:10.1038/mi.2013.31 (2014).

