## Supplemental figures for "Inhibition of IL-22-producing cells by *Sutterella* sp. bacteria"

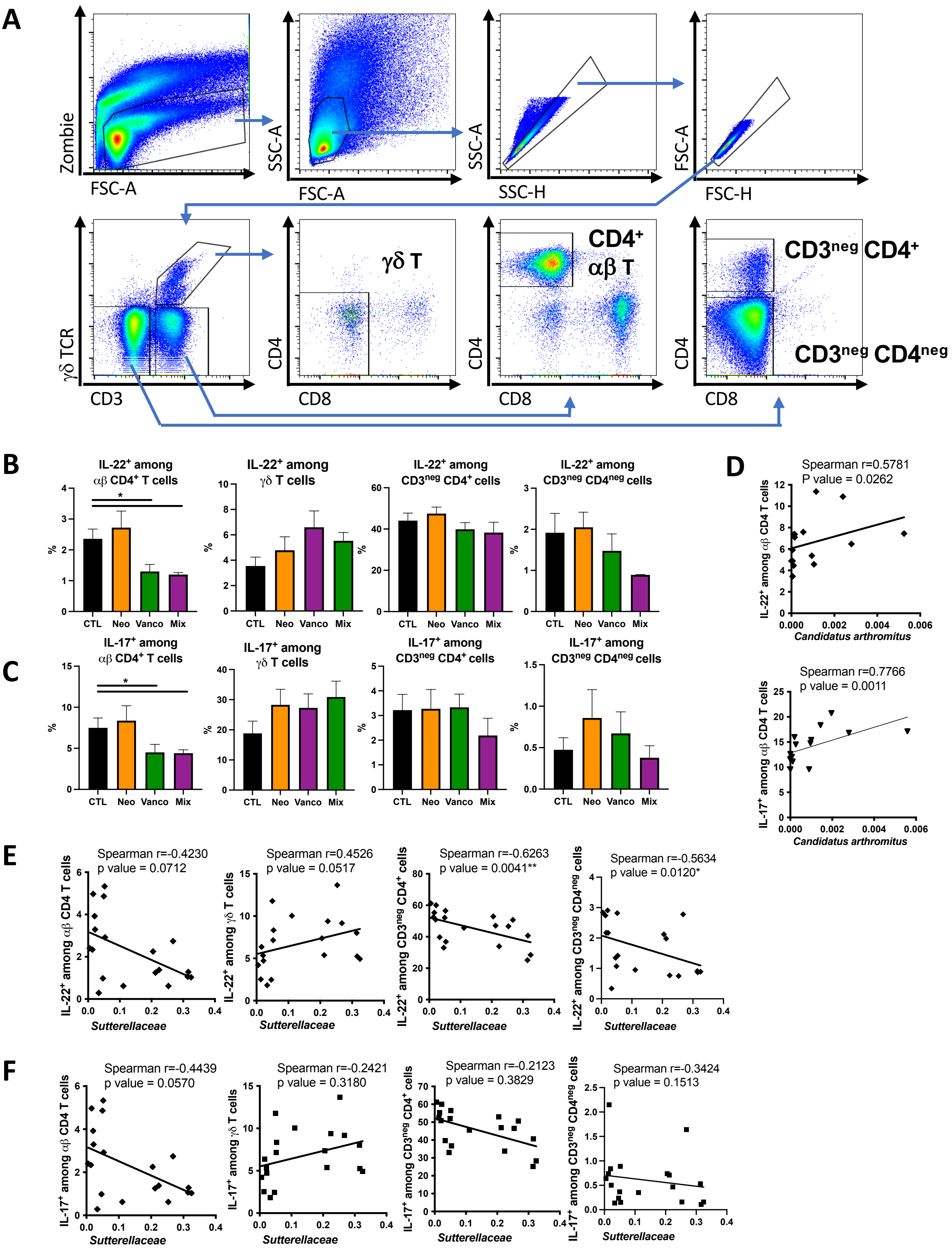


Suppl. Figure 1. ***Sutterella regulates IL-17 and IL-22 productions.*** Related to Figure 1

**(A)** Flow cytometry gating strategy used to identify γδ T cells (γδ TCR$^+$ CD3$^+$ CD4$^{neg}$ CD8$^{neg}$), CD4$^+$ αβ T cells (γδ TCR$^{neg}$ CD3$^+$ CD4$^+$ CD8$^{neg}$), CD4$^+$ CD3$^{neg}$ cells (γδ TCR$^{neg}$ CD3$^{neg}$ CD4$^+$ CD8$^{neg}$) and CD4$^{neg}$ CD3$^{neg}$ cells (γδ TCR$^{neg}$ CD3$^{neg}$ CD4$^{neg}$ CD8$^{neg}$). **(B, C)** Percentage of IL-22 (B) and IL-17 (C) among gated CD4$^+$ αβ T cells, γδ T cells, CD3$^{neg}$ CD4$^+$ and CD3$^{neg}$ CD4$^{neg}$ cells from colon obtained from untreated mice (control, CTL), neomycin (Neo)-, vancomycin (Vanco)– or neomycin+vancomycin (Mix)-treated mice. Cells were stimulated 4 hours with PMA + ionomycin + IL-1β + IL-23. **(D-F)** Correlation by Spearman r between percentage of IL-22 and IL-17-producing CD4$^+$ αβ T cells cells and *C. arthromitus* abondance in small intestine (D) and *Sutterellaceae* abondance in colon (E-F) from mice treated as in B.
Error bars are SEM from 5 to 10 mice per group (B-F). Significant differences were determined using Kruskal-Wallis test, *p < 0.05.

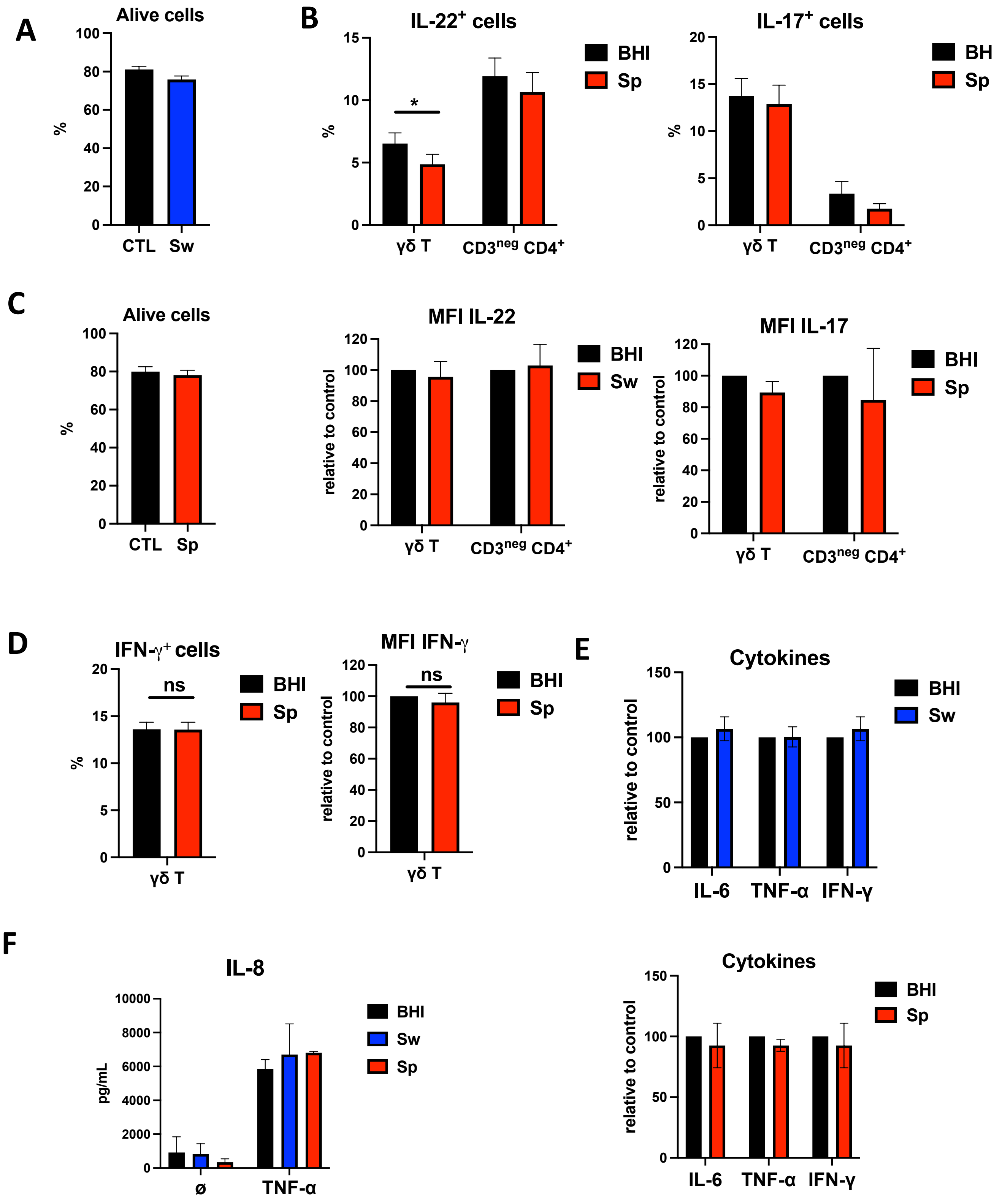


Suppl. Figure 2. ***Sutterella regulates IL-17 and IL-22 productions in vitro.*** Related to Figure 2

**(A)** Proportion of alive cells from pLN cultured 4h with BHI (black) or Sw (blue) culture supernatant and stimulated with PMA + Ionomycin + IL-1β + IL-23 for the 3 last hours. **(B)** Intracellular analysis (top) and normalized geometric mean fluorescence intensity (bottom) of IL-22 (left) and IL-17 (right) expression by gated γδ T cells and CD3$^{neg}$ CD4$^{+}$ cells from pLN cultured with BHI (black), Sp (red) culture supernatant. **(C)** Proportion of alive cells obtained as in B. **(D)** Intracellular analysis (left) and normalized geometric mean fluorescence intensity (right) of IFN-γ expression by gated γδ T cells from pLN cultured with BHI (black), Sp (red) culture supernatant. **(E)** Cells from LP of small intestine were cultured for 24h with BHI (black) or *S. wadworthensis* (Sw, blue) or *S. parvirubra* (Sp, red) culture supernatant and stimulated with PMA + Ionomycin + IL-1β + IL-23. IL-6, TNF-α and IFN-γ productions were measured in the supernatants. **(F)** HT29 cell line was cultured 24h with BHI (black) or *S. wadworthensis* (Sw, blue) or *S. parvirubra* (Sp, red) culture supernatant in presence or absence of TNF-α. IL-8 production was measured in the supernatants.
Error bars are SEM with 13 mice (A), 7 mice (B, C), 4 mice (D), 3-4 mice (E) and 3 biological replicates (F). Significant differences were determined using Wilcoxon test, *P < 0.05.

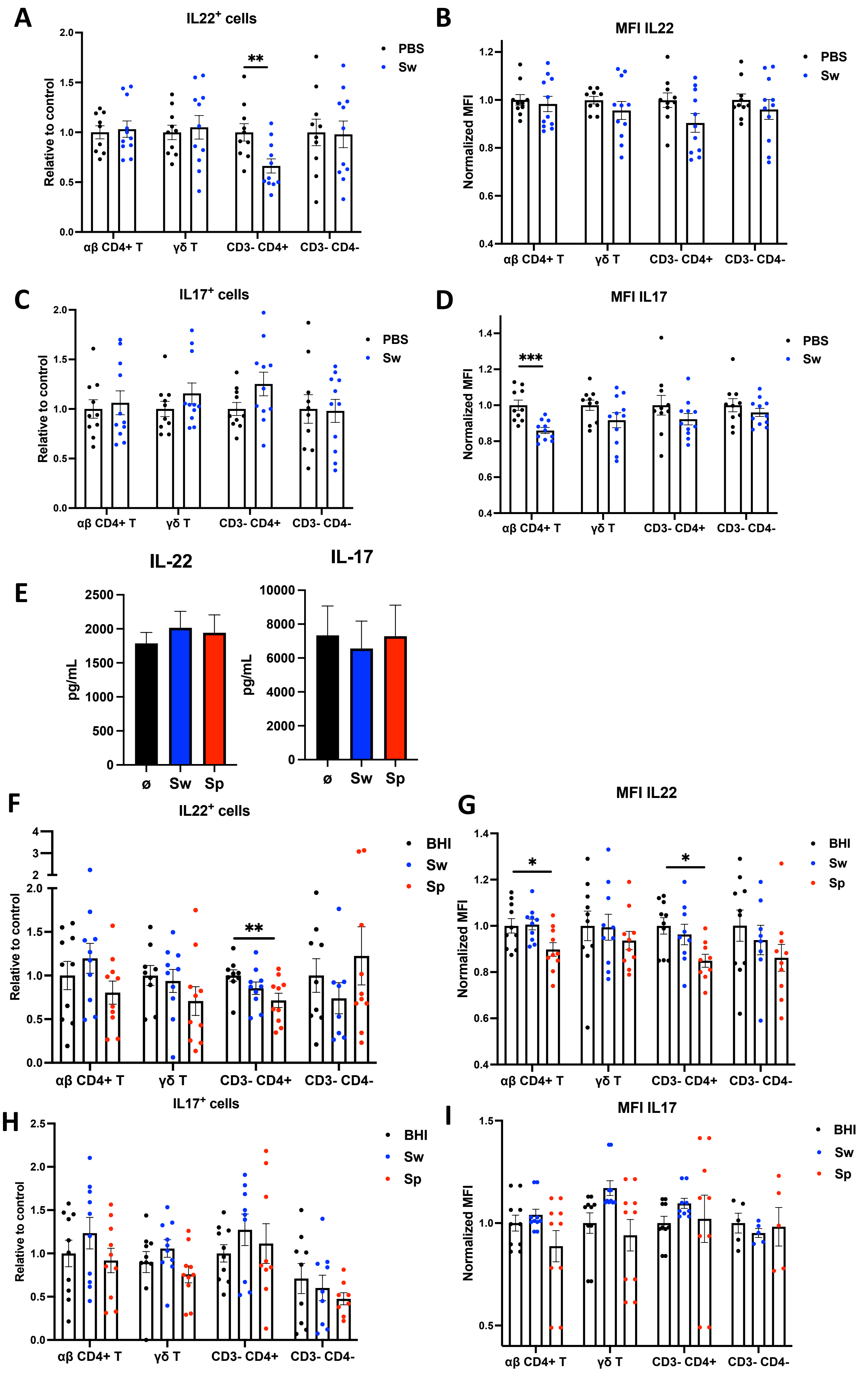
A
IL22+ cells
Relative to control
2.0
1.5
1.0
0.5
0.0
**
PBS
Sw
αβ CD4+ T
γδ T
CD3- CD4+
CD3- CD4-
B
MFI IL22
Normalized MFI
1.2
1.0
0.8
0.6
0.4
PBS
Sw
αβ CD4+ T
γδ T
CD3- CD4+
CD3- CD4-
C
IL17+ cells
Relative to control
2.0
1.5
1.0
0.5
0.0
PBS
Sw
αβ CD4+ T
γδ T
CD3- CD4+
CD3- CD4-
D
MFI IL17
Normalized MFI
1.4
1.2
1.0
0.8
0.6
0.4
***
PBS
Sw
αβ CD4+ T
γδ T
CD3- CD4+
CD3- CD4-
E
IL-22
pg/mL
2500
2000
1500
1000
500
0
ø
Sw
Sp
IL-17
pg/mL
10000
8000
6000
4000
2000
0
ø
Sw
Sp
F
IL22+ cells
Relative to control
4
3
2
2.0
1.5
1.0
0.5
0.0
**
BHI
Sw
Sp
αβ CD4+ T
γδ T
CD3- CD4+
CD3- CD4-
G
MFI IL22
Normalized MFI
1.4
1.2
1.0
0.8
0.6
0.4
*
*
BHI
Sw
Sp
αβ CD4+ T
γδ T
CD3- CD4+
CD3- CD4-
H
IL17+ cells
Relative to control
2.5
2.0
1.5
1.0
0.5
0.0
BHI
Sw
Sp
αβ CD4+ T
γδ T
CD3- CD4+
CD3- CD4-
I
MFI IL17
Normalized MFI
1.5
1.0
0.5
BHI
Sw
Sp
αβ CD4+ T
γδ T
CD3- CD4+
CD3- CD4-

Suppl. Figure 3. ***Sutterella regulates IL-17 and IL-22 productions in vivo.*** Related to Figure 3

**(A-D)** Intracellular analysis of IL-22 (A) and IL-17 (C) expression and normalized geometric mean fluorescence intensity for IL-22 (B) and IL17 (D) by gated $\alpha\beta$ CD4+ T cells, $\gamma\delta$ T cells, $CD3^{neg}$ $CD4^{+}$, $CD3^{neg}$ $CD4^{neg}$ cells from proximal small intestine obtained from PBS (control, black) or *S. wadworthensis* (Sw, blue) -treated mice, and stimulated 3 hours with PMA + Ionomycin + IL-1$\beta$ + IL-23. **(E)** Cells from pLN were cultured for 24h with PBS (black) or dead bacteria (MOI 10:1) (*S. wadsworthensis,* Sw, blue or *S. parvirubra,* Sp, red) and stimulated with PMA + Ionomycin + IL-1$\beta$ + IL-23. IL-22 (left) and IL-17 (right) productions were measured in the supernatants. **(F-I)** Intracellular analysis of IL-22 (F) and IL-17 (H) expression and normalized geometric mean fluorescence intensity for IL-22 (G) and IL17 (I) by gated $\alpha\beta$ CD4+ T cells, $\gamma\delta$ T cells, $CD3^{neg}$ $CD4^{+}$, $CD3^{neg}$ $CD4^{neg}$ cells from proximal small intestine obtained from BHI (left), *S. wadworthensis* (Sw, middle) or *S. parvirubra* (Sp, right) -treated mice, and stimulated as in A.

Error bars are SEM with 9-10 mice per group (A-D, F-I) and 7 mice (E). Significant differences were determined using Wilcoxon test, *$P < 0.05$, **$P < 0.005$ , ***$P < 0.001$.

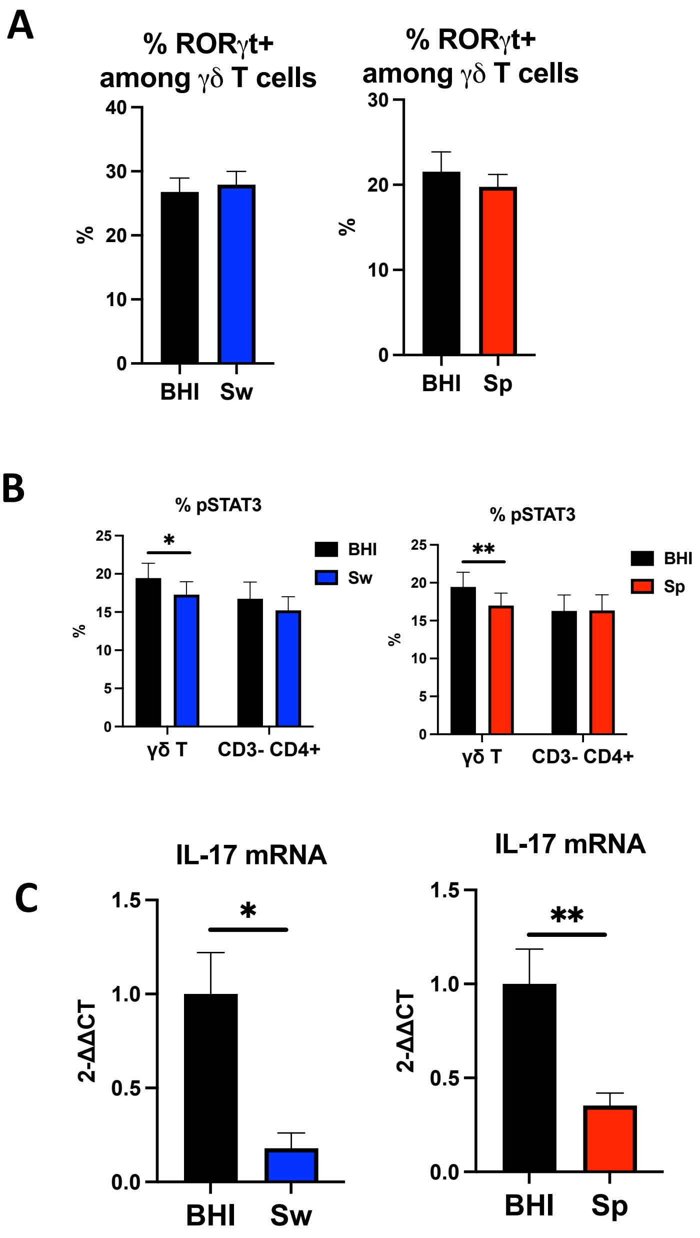


Suppl. Figure 4. ***Sutterella regulates directly IL-22 producing cells.*** Related to Figure 4

**(A)** Proportion of ROR-γt$^+$ cells among γδ T cells from LN cells cultured and stimulated with PMA + Ionomycin + IL-1β + IL-23 for the last 3h **(B)** Proportions of pSTAT3$^+$ cells among γδ T cells from pLN, cultured 2h with BHI or *S. wadsworthensis* (Sw, left) or *S. parvirubra* (Sp, right) culture supernatant and stimulated with IL-23 for the last hour. **(C)** IL-17 mRNA quantification by qPCR in sorted αβ CD4$^+$ T cells cultured 4h with BHI (black) or *S. wadsworthensis* (Sw, blue) or *S. parvirubra* (Sp, red) culture supernatant and stimulated with PMA + Ionomycin + IL-1β + IL-23 for the 3 last hours. Mice were treated with DSS and αβ CD4$^+$ T cells were sorted from mesenteric LN.

Error bars are SEM with 6 mice (A), 8 mice (B), 7-10 mice (C). *P < 0.05, **P < 0.005; by Wilcoxon test.

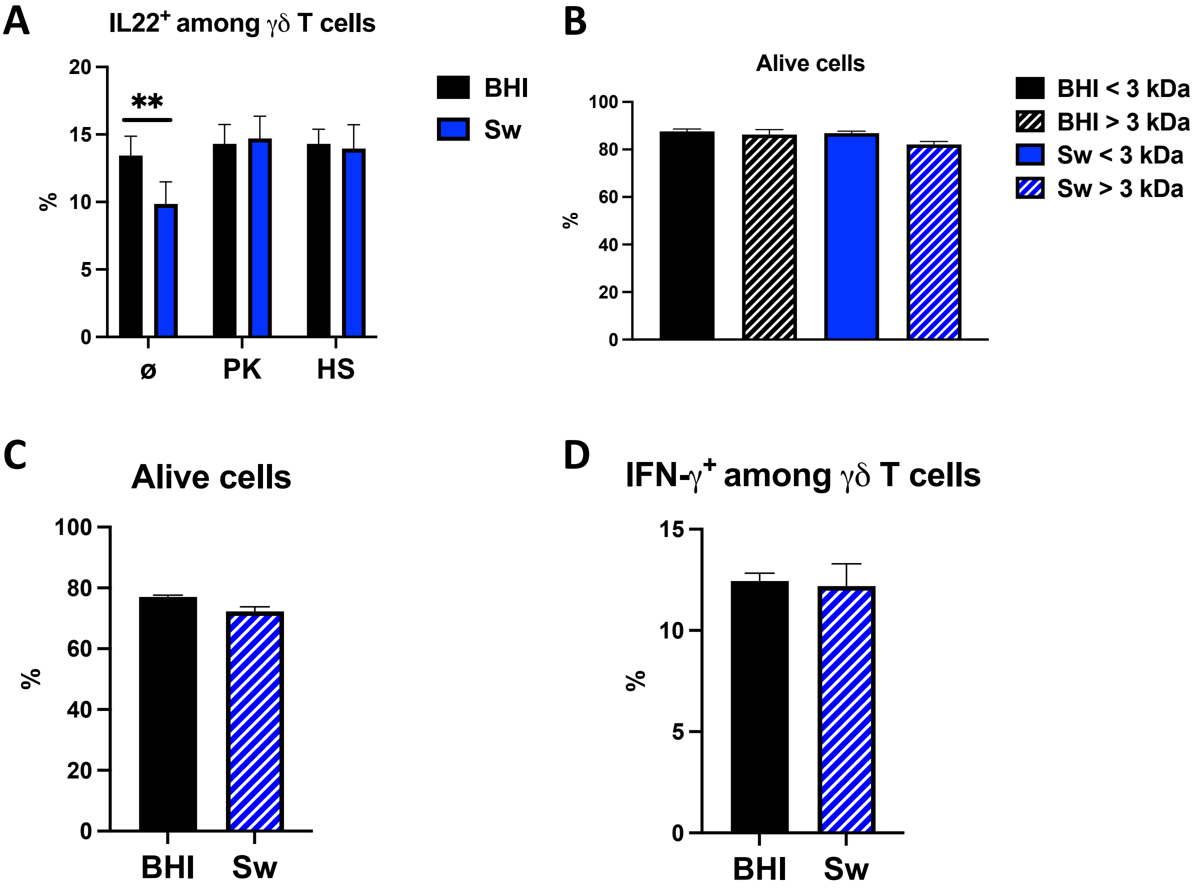


Suppl. Figure 5. **A >30kDa protein from *Sutterella regulates IL-22 productions.*** Related to Figure 5
**(A)** Intracellular analysis of IL-22 expression by gated γδ T cells and CD3$^{neg}$ CD4$^{+}$ cells from pLN cultured 4 hours with BHI or Sw culture supernatant (pretreated with proteinase K (PK) and/or pre-heated (Heat shock, HS) and stimulated with PMA + Ionomycin + IL-1β + IL-23 for the 3 last hours. **(B-C)** Proportion of alive cells from pLN cultured 4 hours with <3 kDa and >3 kDa fractions (B) or > 30kDa (C) of BHI or Sw culture supernatant and stimulated as in A. **(D)** Intracellular analysis of IFN-γ expression by gated γδ T cells cultured and stimulated as in C.
Error bars are SEM from 6 mice (A), 7 mice (B) and 5 mice (C, D). Significant differences were determined using paired t test (A) and Wilcoxon test (B-D): *P < 0.05, **P < 0.005 , ****P < 0.0001. See also Figure S5
